# Predicting phenotypes with one step genetic decision trees

**DOI:** 10.1101/2025.05.29.656727

**Authors:** Julie Blommaert, Philipp E. Bayer, David Ashton, Georgia Samuels, Linley Jesson, Maren Wellenreuther

## Abstract

Genomic prediction of complex traits is limited when phenotype records are restricted and when using linear models. Increasing the amount of phenotypic data with high-throughput, image-based phenotyping could result in better genomic prediction and stronger signals in variant detection. Here, we analysed phenotypic and genomic data from a selectively bred cohort of the Australasian snapper (*Chrysophrys auratus*) to identify genetic variants associated with growth traits. We used a high-throughput phenotyping pipeline to extract 13 measurements of size from images. Phenotypic correlations among image-derived and manually measured traits (weight, fork length), together with heritabilities, were analysed. All measurements were significantly positively correlated with each other, and heritability ranged from 0.20-0.38. Genome-wide association studies (GWAS) identified 28 growth-associated SNPs, while GBLUP was used to predict phenotypes, and XGBoost machine-learning models were used to jointly predict phenotypes and report important variants. Both GBLUP (mean R^2^ = 0.50) and XGBoost (mean R^2^ = 0.77) performed well on the training data, but performance dropped on testing sets (both = 0.11), which decreased further when accounting for genetic relatedness (both = 0.06). Despite this, approximately 20% of the genetic variance for growth traits was captured by the models, and feature importance from XGBoost reflected signals seen in GWAS. Our findings highlight the utility of integrating computer vision-based phenotyping with GWAS, GBLUP, and ML for trait prediction. Despite detecting shared biological signals as GWAS, genomic prediction faces challenges with population structure and relatedness that are inherent in breeding programmes of mass spawning species, including many aquatic species.

**Article summary:** Incorporating genetic information into selective breeding programmes can accelerate gains but may also miss gene interactions in complex traits. Machine learning approaches, such as decision trees, can capture those relationships and potentially improve genomic predictions. We used high-throughput computer vision phenotyping to uncover biological signal for genetic growth variants involved in the Australasian snapper. Both types of models captured 18-40% of the genetic variation, but the prediction accuracies were hampered by the population structure in this cohort of mass-spawning fish.

## Introduction

Selective breeding programmes are fundamental to both plant and animal production systems, where they are used to improve economically important traits such as growth, yield, survival, product quality, reproductive performance, and feed efficiency (Godfray et al. 2010; Boyd et al. 2022). Over recent decades, the increasing availability of genomic resources, including whole-genome assemblies and dense panels of DNA markers, has transformed selective breeding approaches, even in non-model species (Ekblom and Galindo 2011; Ellegren 2014). Genetic improvement has progressively shifted from conventional pedigree-based selection towards marker-assisted and genomic selection, or combinations of both approaches (Sandoval-Castillo et al. 2022). These advances have substantially improved the estimation of breeding values and the identification of superior breeders without the need for additional phenotyping, accelerating genetic gain across a wide range of production systems. This is especially important where mass phenotyping is not practical.

Despite these advances in genomics, phenotyping remains a major bottleneck limiting further progress in breeding programmes (Houle 2010). While genomic data generation has become increasingly affordable and scalable, the collection of accurate, high-throughput phenotypic data remains challenging for many economically important and resilience-related traits. This challenge is particularly pronounced for traits that are labour-intensive, invasive, expensive, or difficult to measure repeatedly across large numbers of individuals. In aquaculture systems, phenotyping challenges are especially severe because of the aquatic environment itself (Fu and Yuna 2022). Fish must often be removed from water, anaesthetised and handled before measurements can be obtained, creating substantial logistical constraints and introducing handling stress that can affect animal welfare and, in some cases, phenotype expression. Importantly, many measurements must be completed within very short time windows while fish remain under anaesthesia, limiting the number and complexity of traits that can realistically be collected manually. This makes the measurement of traits such as morphology, behaviour, stress responses, welfare indicators, and condition particularly difficult at large scales. Consequently, phenotyping has emerged as one of the key constraints limiting the efficiency and scalability of aquaculture breeding programmes.

Recent technological advances in machine learning (ML), computer vision, automated imaging, and deep learning networks provide new opportunities to overcome these limitations (Yang et al. 2021). Image-based computer vision approaches have transformed phenotyping in aquaculture by enabling rapid, non-invasive, and high-throughput trait extraction from images and videos (Tuckey et al. 2022; Babu et al. 2023). These approaches substantially reduce handling time, minimise stress on animals, and allow the capture of detailed morphological and behavioural information that would otherwise be difficult or impossible to measure manually. In aquaculture, computer vision systems have increasingly been used to estimate growth, body shape, swimming behaviour, health status, feeding activity, deformities, and biomass in species such as salmon, seabream, tilapia, shrimp, and other cultured fish species. Deep learning frameworks, particularly convolutional neural networks (CNNs), have shown strong performance in automated trait extraction and classification tasks, substantially reducing manual labour and observer bias.

At the same time, ML approaches are increasingly being applied to genomic prediction and variant discovery (Chafai et al. 2023; Song et al. 2023; Yáñez et al. 2023; Chen et al. 2025). Traditionally, breeding decisions integrate phenotypic and pedigree or genomic information through best linear unbiased prediction (BLUP) frameworks, including genomic BLUP (GBLUP), to estimate breeding values. Where the underlying genetic architecture of traits is of interest, genome-wide association studies (GWAS) are commonly employed to identify trait-associated loci. However, these approaches can become computationally demanding and less powerful in emerging breeding programmes with limited sample sizes, sparse genomic resources, or highly polygenic traits. Machine learning approaches may provide complementary solutions to these challenges (Zhao et al. 2021). Decision-tree based methods such as Random Forests and Extreme Gradient Boosting (XGBoost) can simultaneously perform phenotype prediction and genotypic variant prioritisation by identifying genomic regions with high predictive importance. Unlike traditional GWAS approaches that test markers individually, these methods can capture non-linear interactions, epistasis, and complex genotype–phenotype relationships. In livestock, plant and aquaculture systems, ML approaches have already been explored for predicting disease resistance, growth performance, feed efficiency, and environmental tolerance traits (Okser et al. 2014; Bassim et al. 2015; He and Knowles 2016; Chen et al. 2025). Importantly, these methods may also facilitate more targeted future genotyping strategies by identifying smaller subsets of highly informative markers for breeding applications.

The Australasian snapper (Sparidae: *Chrysophrys auratus*), also known in New Zealand as tāmure, represents a promising candidate species for warm water aquaculture diversification in New Zealand. Snapper has a broad latitudinal distribution across northern New Zealand and southern Australia and naturally experiences a wide range of thermal environments, from temperate southern waters below 10°C to subtropical northern habitats reaching temperatures of up to 30°C, and individuals generally display a broad tolerance to these temperatures (Bentley-Hewitt et al. 2024; Bentley-Hewitt et al. 2025). This broad thermal tolerance suggests considerable potential for farming under diverse environmental conditions and future climate scenarios. Over the past two decades, substantial advances have been made in developing snapper aquaculture in New Zealand, including improvements in husbandry, production cycles, and selective breeding (Moran et al. 2023; Samuels et al. 2024; Samuels et al. 2026). Recent breeding programmes have established elite snapper lines with improved growth, survival, and feed conversion efficiency (Samuels et al. 2024; Samuels et al. 2026), alongside the development of genomic resources and breeding tools (Montanari et al. 2023; Bentley-Hewitt et al. 2024; Blommaert et al. 2024).

Here, we integrate high-throughput image-based phenotyping, genomic analyses, and machine learning approaches within a selective breeding framework for snapper. Specifically, we 1) perform computer vision-based phenotyping to extract growth-related traits, 2) apply traditional genomic prediction (GBLUP) and genome-wide association studies (GWAS) to estimate breeding values and identify trait-associated variants, and 3) combine genomic prediction and variant discovery within a single machine-learning framework using decision-tree approaches implemented in XGBoost. By integrating phenomics, genomics, and ML-based prediction, this study evaluates how data science approaches can accelerate aquaculture breeding programmes and support the development of climate-resilient and robust aquaculture species enabling future work on genomic selection to traits that are more difficult to phenotype (e.g. physiological responses, commercial traits such as fillet size and quality, or individual feed conversion ratios).

## Materials and Methods

### Fish rearing

Snapper used in this study came from the long-term selective breeding programme at The New Zealand Bioeconomy Science Institute Ltd. (BSI), Nelson, New Zealand. Genetic and growth data were obtained from the F_4_ generation, produced in November 2021 from the third selectively bred generation. Fertilized eggs were collected and incubated for 5 days. Larval rearing protocols followed standard methods developed for this species (Samuels et al. 2024). Fish were then reared at BSI’s land-based fish facility. The facility uses a flow-through tank system, where water from the Nelson Haven is withdrawn from the intertidal zone via an engineered aquifer filled with various hard substrates that provide filtration. All work was done under the conditions of animal ethics application AEC-2021-PFR-05, approved by the Animal Ethics Committee at the Nelson Marlborough Institute of Technology (Te Pūkenga), along with fish-farm licence FW208.03.

### Manual and image-based phenotyping

Prior to any measurements, snapper were anaesthetised with AQUI-S® (15–20 ppm), sufficient to induce loss of equilibrium and absence of response to handling. At three months of age, fish were weighed (FX-300i WP, A&D Company Ltd.), photographed and measured manually. Additional phenotypic measurements (Fig. 1A) were obtained from the photographs using an computer vision pipeline that extracts traits such as length, and specific body area measurements, directly from the image. Measurements from this pipeline are available in Supplementary Table 1. In total, 1011 individual fish were phenotyped by imaging. The first principal component (PC) derived from measured traits (using prcomp from the stats library in R) were included in downstream analyses. Trait correlations were assessed using the R package corrplot v0.92.

**Figure 1.**
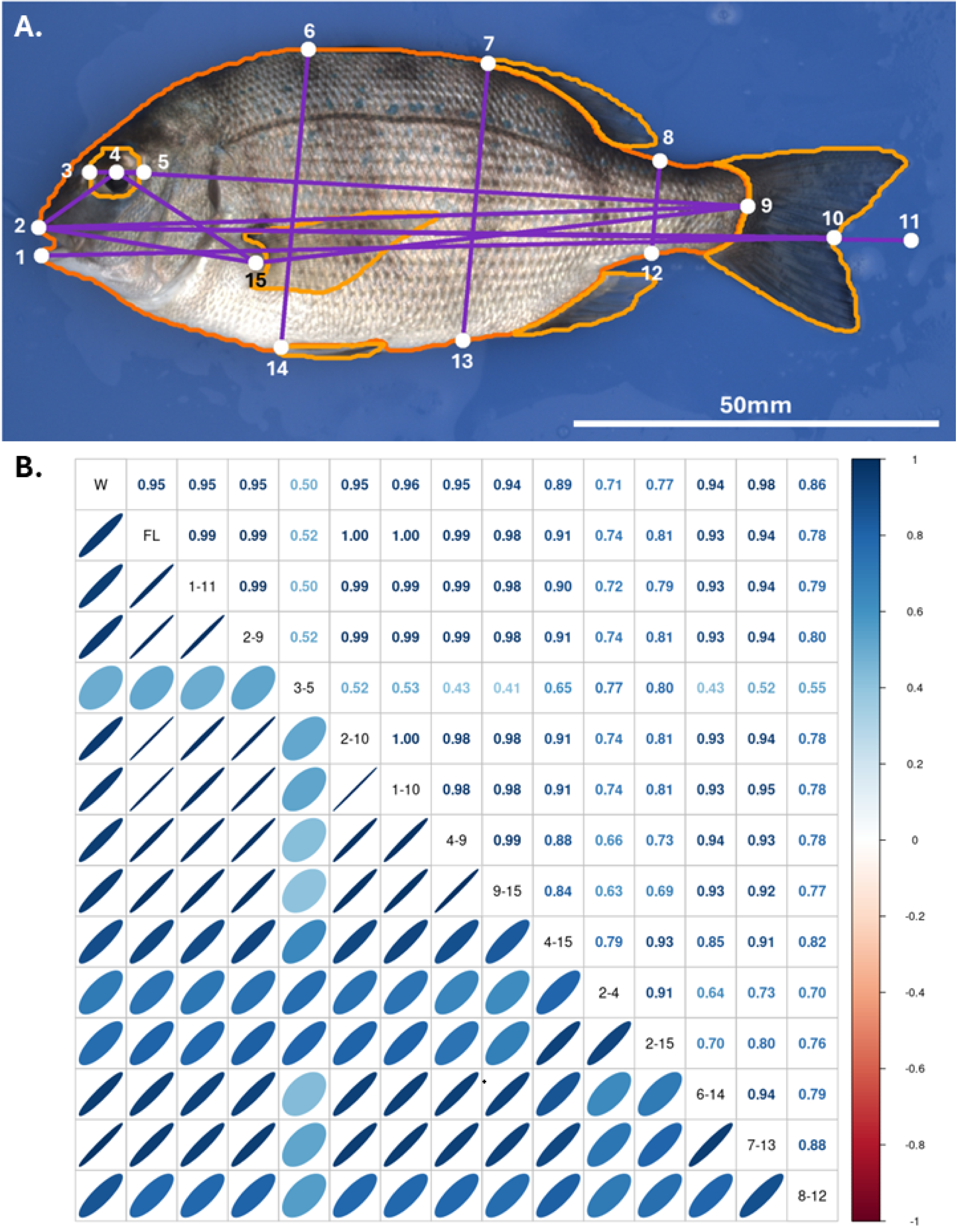
**A)** An example output of the computer vision phenotyping. The contours of fish body parts (orange) and landmarks (white points) used for each of the 13 measurements gathered by this pipeline (purple). The white line indicates a scale bar of 50mm. The labelled points indicate landmarks used for measurements from the computer vision pipeline. Landmarks are 1-bottom lip, 2-top lip, 3 and 5-the left and right edges of the eye respectively, 4-centre of the eye, 6 and 14-top and bottom of the fish at 25% of its total length respectively, 7 and 13-top and bottom of the fish at 50% of its total length respectively, 8 and 12-top and bottom of the fish at 75% of its total length respectively, 9-peduncle, 10-tail fork, 11-total length end point. **B)** Correlation matrix of all 15 measured traits in this study. The colour and shape of each ellipse (bottom half) represents the R^2^ value (top half) for that correlation (written in the corresponding box for each correlation). R^2^ values were only included where p-values were < 0.05.

### Genetic data collection

SNP data for these fish were collected using a multi-species SNP chip (Montanari et al. 2023). This SNP chip included 18,489 snapper SNPs, but after filtering for quality and levels of polymorphism 11,164 SNPs were retained for the 1,011 fish included in this work. SNP positions were remapped from the original snapper reference genome to the updated snapper genome assembly (Blommaert et al. 2024) using snplift v1.0.4 (Normandeau et al. 2023). SNPs that could not be lifted over were further processed by extracting a genomic region of 70 bp surrounding each SNP from the original assembly and performing a BLAST search against the new genome assembly. SNPs with multiple mapping locations in the updated genome were excluded, as were SNPs that did not align to the expected chromosome.

### Genetic relatedness and heritability

Genetic relatedness among individuals was estimated from SNP genotype data to visualise population structure within the F_4_ fish and to stratify the data set for genomic prediction analyses. Pairwise relatedness coefficients were calculated in R by centering the genotype matrix (subtracting the column-wise mean genotype value for each SNP) and computing the genomic relationship matrix (GRM) as the dot product of the centred matrix and its transpose, divided by the total number of SNPs. We grouped genetically similar individuals using k-means clustering (k = 19) applied to the GRM. The resulting cluster assignments were visualised by principal component analysis (PCA) to confirm consistency between the k-means groupings and the genetic structure apparent in the PCA.

We estimated narrow-sense heritability for each trait using ASREML-R v4.2 (Bulter et al. 2023), fitting mixed-effects models with fish id as a random effect. Variance components were estimated by restricted maximum likelihood (REML) and heritability was calculated as the proportion of total phenotypic variance explained by additive genetic effects. Standard errors were obtained using the *predict* function in ASREML.

### Effective population size (N*e*) estimation

We estimated N*_e_* based on the level of LD only between all markers that are on different chromosomes in the genome (i.e. interchromosomal LD). This approach eliminates the confounding effects of physical linkage and recombination hotspots. To ensure high-quality markers for linkage disequilibrium (LD) analysis, we filtered the SNP chip data based on minor allele frequency (MAF) using PLINK v1.90b6.5 (Purcell et al. 2007). Variants were excluded if the MAF was below 0.05. Additionally, a more conservative MAF filter of 0.1 was also applied. Interchromosomal *r^2^* was calculated for all possible SNP pairs across the 24 chromosomes using PLINK and N*_e_* was then estimated from this using the bias correction method (Waples 2006):

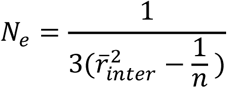

where 1/*n* represents the expected *r^2^* due to sampling error alone. This approach provided a measure of the background genetic structure and shed some light on the reproductive skew within the F_4_ lineage of the breeding programme.

### Identifying growth-related variants using GWAS

We conducted genome-wide association studied (GWAS) using rMVP v1.1.1 (Yin et al. 2021), applying three standard models: general linear model (GLM), mixed linear model (MLM, and Fixed and Random model Circulating Probability Unification (FarmCPU). For each trait, we modelled the phenotype as a function of the SNP being tested, with the first two genomic principal components included as covariates to account for population structure. The MLM additionally accounted for relatedness among individuals by fitting a random polygenic effect based on the genomic relationship matrix (GRM). FarmCPU iteratively fit SNPs as the fixed effects while controlling for confounding factors using a random effect, improving power and reducing false positives when compared with GLM and MLM.

We performed these analyses for all 15 measured traits as well as condition factor (K) and the phenotypic PC1 (explaining 97.16% of phenotypic variability). Significance thresholds were determined using Bonferroni correction (α = 0.05), and Manhattan and quantile-quantile (QQ) plots were inspected to assess model inflation. Downstream comparisons and functional analyses focused on SNPs identified by the FarmCPU model.

SNP functional annotation was performed via snpEff v5.2 (Cingolani et al. 2012) using the genome assembly and snapper specific annotations from previous work (Blommaert et al. 2024). Functional profiling was performed with g:GOSt on the g:profiler website (Kolberg et al. 2023). Other functional information was gathered from the Zebrafish Information Network (ZFIN) (Bradford et al. 2022).

### Cross validation of genomic prediction using GBLUP

We used GBLUP to predict weight, length, PC1 of the phenotypes, eye width, and height at 75% of the fish’s length from genome-wide SNP data using ASREML v4.2. We chose these traits to represent the range of heritabilities in our dataset and to investigate if there was any increased signal in composite traits (e.g. PC1) compared to individual measurements. The previously described GRM was used, with a small value added to the diagonal prior to computing its inverse to ensure numerical stability. We evaluated two prediction strategies to assess the impact of relatedness between training and testing individuals.

In both approaches, we portioned individuals into training and testing sets and individuals in the testing set were masked by setting their measured phenotype values to missing. In the first approach, individuals were randomly partitioned (80:20, train:test), and in the second, entire relatedness clusters were assigned to training (80% of clusters) and test (20% of clusters). The cluster-based splits aimed to reduce genetic relatedness between sets. To obtain robust estimates of prediction, we repeated both partitioning strategies 50 times using different randomisation seeds, and prediction accuracy metrics compared to the real measurement were reported for each of these replicates.

For both approaches, we fitted mixed-effects models with an intercept as the only fixed effect. Random effects included an additive genetic effect for each individual, modelled using the inverse GRM, and cluster membership was fitted as an independent random effect to account for residual population structure. We estimated variance components using restricted maximum likelihood.

### Nested cross-validation for genomic prediction using machine learning

Genomic prediction analyses for weight, length, PC1 of the phenotypes, eye width, and height at 75% of the fish’s length were performed using the *XGBoost* (Chen and Guestrin 2016) algorithm in the R package xgboost v1.7.8.1. Models were tuned using tidymodels v1.2.0 (Kuhn and Wickham 2020) with 5-fold internal cross-validation. We evaluated performance using root mean square error (rmse) between predicted values and real measurements as a metric. To minimise potential data leakage due to genetic relatedness, we partitioned the dataset into training (80%) and testing (20%) subsets 1) randomly, and 2) based on the relatedness clusters identified above, as was done for GBLUP. As above, these CV splits were performed 50 times, using the same seeds as for the GBLUP cross validation.

Model preprocessing was defined using a *recipes* workflow. We included all SNP genotypes as predictors, while individual identifiers were retained only as sample identifiers and excluded from model fitting. To account for genetic structure and reduce dimensionality, we applied principal component analysis (PCA) to numeric predictors, with the number of retained components treated as a tunable hyperparameter. The original SNP variables were retained alongside PCA components to allow the model to use both raw genotypes and derived structure information. Genetic relatedness clusters were included as categorical predictors and encoded using one-hot (dummy) variables.

XGBoost models were fitted in regression mode using root mean square error (RMSE) as the optimisation metric. We specified a maximum of 500 trees, with early stopping applied to halt training when validation performance failed to improve for 20 consecutive iterations. The following hyperparameters were tuned: tree depth, minimum number of observations per node, loss reduction (gamma), number of variables sampled at each split (mtry), learning rate, and the number of PCA components. Hyperparameter tuning was performed using a Latin hypercube sampling strategy with 20 candidate parameter combinations. Hyperparameter search ranges are provided in Table 1. We assessed model performance using internal 5-fold cross-validation, with folds grouped by genetic relatedness cluster to prevent information leakage between training and validation sets. The optimal hyperparameter combination was selected based on a stability criterion that balanced low mean RMSE with low variance in RMSE across cross-validation folds. We then refitted the final model to the full training dataset using the selected parameters.

**Table 1.** Hyperparameter search ranges explored in XGBoost tuning.

| Parameter | Range explored |
| --- | --- |
| Tree depth | 2-10 |
| Min n | 2-50 |
| Mtry | 2-200 |
| Learning rate | 0.001-0.1 |
| Loss reduction | 0-1 |
| PCA components | 2-6 |

Predictive performance was evaluated by generating predictions for both the training and testing sets. Model accuracy was assessed using RMSE and the coefficient of determination (R²). We retained importance matrices from the final XGBoost models for downstream analyses.

We also extracted the feature importance matrix from each of the 50 cross validation replicates and produced a summary table including mean gain, mean cover, and a count table for how often each SNP occurred in an importance matrix across the 50 cross-validation runs for either the cluster based split or the individual based splits. To evaluate the feature selection reliability, SNPs were grouped by the number of CVs they were selected in (out of 50). SNPs that were selected in 35 CVs or more were classed as persistent markers, and those that were selected in fewer than 35 CVs were designated as background markers. We used mean gain across the 50 CVs as the primary measure of feature importance. A non-parametric bootstrapping approach in R was taken to compare the mean gain between the persistent and background markers. For each stratification approach and SNP stability category, the observed pool mean of SNP mean gain values was resampled over 1000 independent bootstrap iterations. The bootstrap sample mean was calculated for each iteration and unified 95% confidence intervals (CIs) were extracted. To test the directional hypothesis that persistent markers exhibit significantly higher feature importance than transient background noise, a one-tailed Mann-Whitney *U* (Wilcoxon rank-sum) test was applied to the empirical distributions.

## Results

### Phenotypic and genotypic variation

Summary statistics for each of the 15 measured phenotypes across the 1011 fish can be found in Table 2. Weight ranged from 7.46g to 45.19g, while fork length ranged from 74.47mm to 123.03mm. Overall, traits were highly positively correlated with each other (Fig. 1B), and 97.16% of the phenotypic variance was explained by PC1 of the phenotypes. Correlation coefficients ranged from 0.41 (eye width vs. distance between the caudal peduncle and pectoral joint) to 1 (fork length versus distances between each lip and the tail fork, and distance between top lip and tail and distance between bottom lip and tail fork) (Fig. 1B).

**Table 2.** Summary of phenotypic measurements and heritability estimates. Measurements are reported as minimum, mean, and maximum values, with corresponding narrow-sense heritability estimates (h²). The standard error of all h^2^ estimates was 0.05 when rounded to two decimal places. Landmark numbers indicate the points on Figure 1 used to derive each linear measurement; “–” indicates direct measurements not derived from landmarks.

| Measurement | Landmarks | Minimum | Mean | Maximum | $h^2$ |
| --- | --- | --- | --- | --- | --- |
| weight_g | - | 7.46 | 25.46 | 45.19 | 0.38 |
| fork_length_mm | - | 74.47 | 104.68 | 123.03 | 0.33 |
| total_length_mm | 1-11 | 80.10 | 112.61 | 132.73 | 0.34 |
| standard_length_mm | 2-9 | 66.93 | 95.31 | 112.27 | 0.33 |
| eye_width_mm | 3-5 | 5.83 | 7.66 | 9.16 | 0.20 |
| top_lip_tail_fork_mm | 2-10 | 73.50 | 103.60 | 121.70 | 0.34 |
| bottom_lip_tail_fork_mm | 1-10 | 73.40 | 103.55 | 122.03 | 0.34 |
| eye_caudal_peduncle_mm | 4-9 | 53.97 | 79.75 | 94.50 | 0.33 |
| caudal_peduncle_pectoral_joint_mm | 9-15 | 41.03 | 62.83 | 75.50 | 0.33 |
| eye_pectoral_joint_mm | 4-15 | 15.70 | 21.14 | 25.37 | 0.31 |
| eye_top_lip_mm | 2-4 | 8.87 | 12.04 | 15.77 | 0.24 |
| pectoral_joint_top_lip_mm | 2-15 | 20.83 | 27.60 | 33.87 | 0.25 |
| height_0.25_mm | 6-14 | 23.70 | 36.68 | 46.10 | 0.33 |
| height_0.5_mm | 7-13 | 25.53 | 37.56 | 46.77 | 0.38 |
| height_0.75_mm | 8-12 | 9.13 | 13.72 | 24.85 | 0.34 |

In total, after QC and SNP lift over, 158 SNPs were not transferred to the new assembly, and 11,006 SNPs were included in downstream analyses. Genetic relatedness corresponded to the clusters on the PCA plot, and PC1 explained 7.55% of the variance in the SNPs, while PC2 explained 5.63% (Fig. 2A). In the absence of a pedigree, we used the GRM and k-means clustering to partition the individuals into 19 clusters, which corresponded well to the clusters on the PCA plot (Figure 2A). Of these clusters, four were significantly lighter (p < 0.05) than the overall mean (25.47 g), and four significantly heavier (Figure 2B). Estimated heritabilities ranged from 0.201 (eye_width_mm) to 0.3792 (height_0.5_mm). Notably, the estimated heritability of weight was also moderately high at 0.3775 (Table 2).

**Figure 2.**
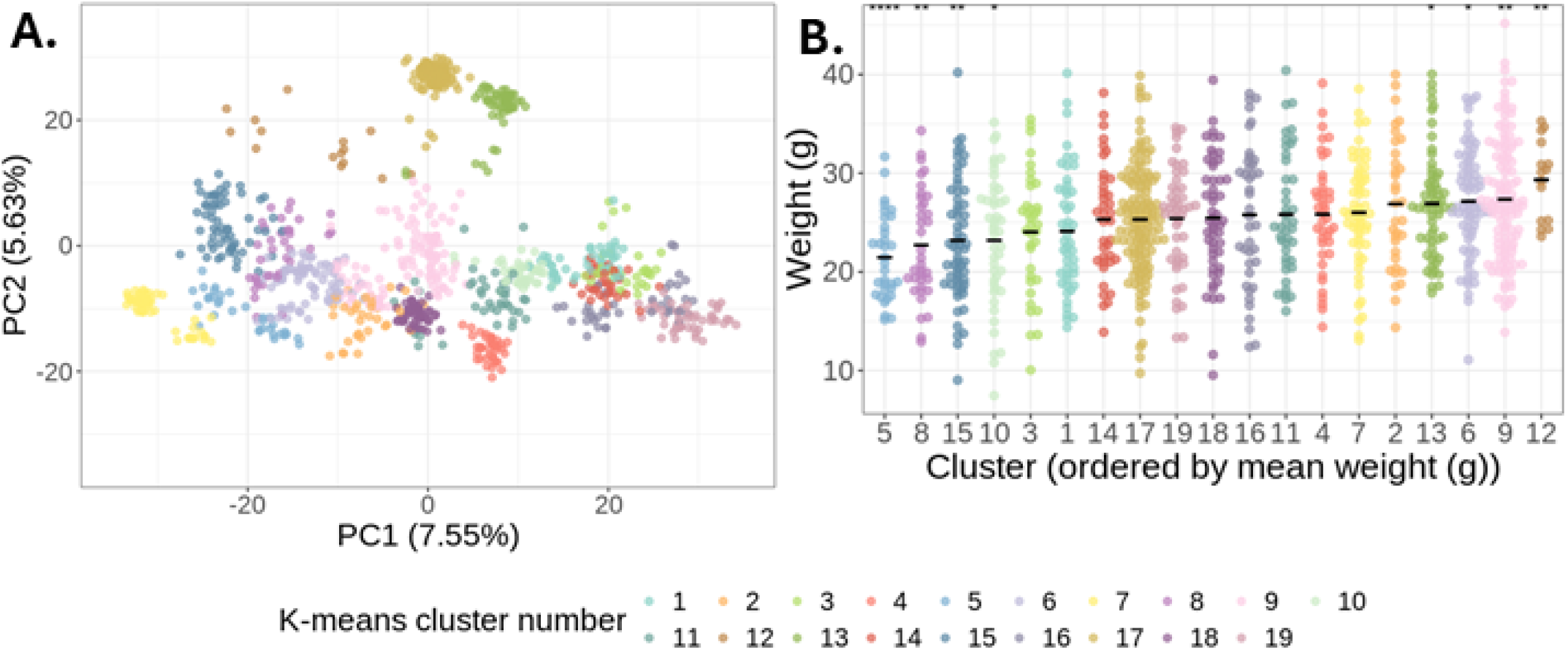
**A)** Principal component (PC) clustering based on the SNP chip genotypes. Each point represents an individual fish (n = 1011). Total explained variance of each PC is shown in brackets on the axis label for that PC. Clusters are coloured based on relatedness clusters determined by *k-means* clustering (k = 19) of the genetic relatedness matrix. **B)** Weight (in g) of the fish grouped by relatedness cluster. Clusters are ordered on the x-axis by mean weight, which is indicated with a black bar for each cluster. Significant differences in weight between clusters and the overall average weight is indicated by asterisk (* p_value < 0.05, ** p_value < 0.01, *** p_value < 0.001).

The mean r^2^ of the interchromosomal linkage disequilibrium (LD) was between 0.0186 (MAF > 0.05) and 0.0193 (MAF > 0.1), but this was skewed much higher than the median (0.0071-0.0068). This skew was driven by a small number of SNPs with an LD r^2^ < 0.8 in each case (148-158) so both values were used in the Waples equation to estimate the N_e_. When using the mean r^2^ the N_e_ estimate was 18.2-18.9, and when using the median, it was 54.7-57.8 (Table 3).

**Table 3.**
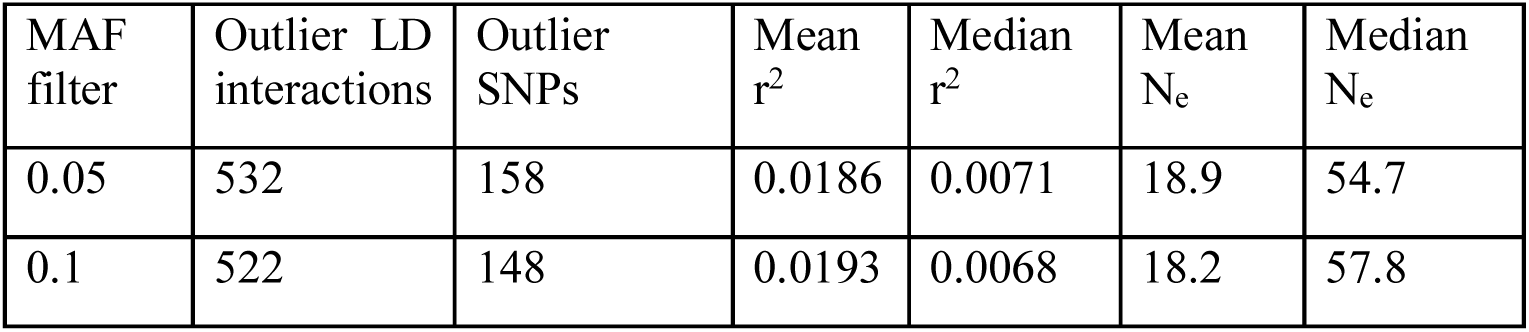
For each MAF filter, the number of outlier interchromosomal LD interactions (r^2^ < 0.8), the number of unique SNPs involved in these interactions, the mean and median r^2^ for the interchromosomal LD interactions, and the estimated N_e_ calculated using the mean or the median.

| MAF filter | Outlier LD interactions | Outlier SNPs | Mean $r^2$ | Median $r^2$ | Mean $N_e$ | Median $N_e$ |
| --- | --- | --- | --- | --- | --- | --- |
| 0.05 | 532 | 158 | 0.0186 | 0.0071 | 18.9 | 54.7 |
| 0.1 | 522 | 148 | 0.0193 | 0.0068 | 18.2 | 57.8 |

### Identifying growth-related SNPs using GWAS

Of the three GWAS methods included in rMVP, MLM identified no associated SNPs with any phenotypes, GLM showed p-value inflation for almost all p-values (Figure 3, Supplementary Figures 1 and 2), and the number of SNPs identified by FarmCPU (Supplementary Table 2) as being significantly associated with a trait ranged from two (caudal_peduncle_pectoral_joint_mm) to eight (eye_top_lip_mm and height_0.75_mm). For PC1 of the phenotype, seven significant SNPs were identified using FarmCPU. Across all phenotypes with FarmCPU (including phenotypic PC1), 29 SNPs were found to be associated, with 20 being unique to the trait they were identified in, and nine shared across at least two phenotypes (Supplementary Table 3). Significant SNPs were distributed across 16 of the 24 chromosomes in the snapper genome with the highest number of SNPs (four) on chromosomes 3 and 5. The 29 SNPs were found to affect 14 annotated genes with modifier effects (Supplementary Table 3). Of these genes, twelve (arrdc1a, clstn2a, mybpha, osbpl3b, pde4dip, tpst1, zdhhc3a, atp5pf, eya4, ptprub, tln2a, pitpnb) were found to be targets of the *Hox9a:meis1a* transcription factor complex, four (osbpl3b, pde4dip, zdhhc3a, leap2) were found to be targets of transcription factor *p53*, and six (clstn2a, osbpl3b, pde4dip, soat1, tpst1, zdhhc3a) were found to be targets of the transcription factor *Sox-10*. The two SNPs that were significant across the most phenotypes were found to affect genes interacting with *Sox-10*. The other genes affected by the significant SNPs were a glutamate receptor (grik5) and a potassium voltage-gated channel (kcna7).

**Figure 3.**
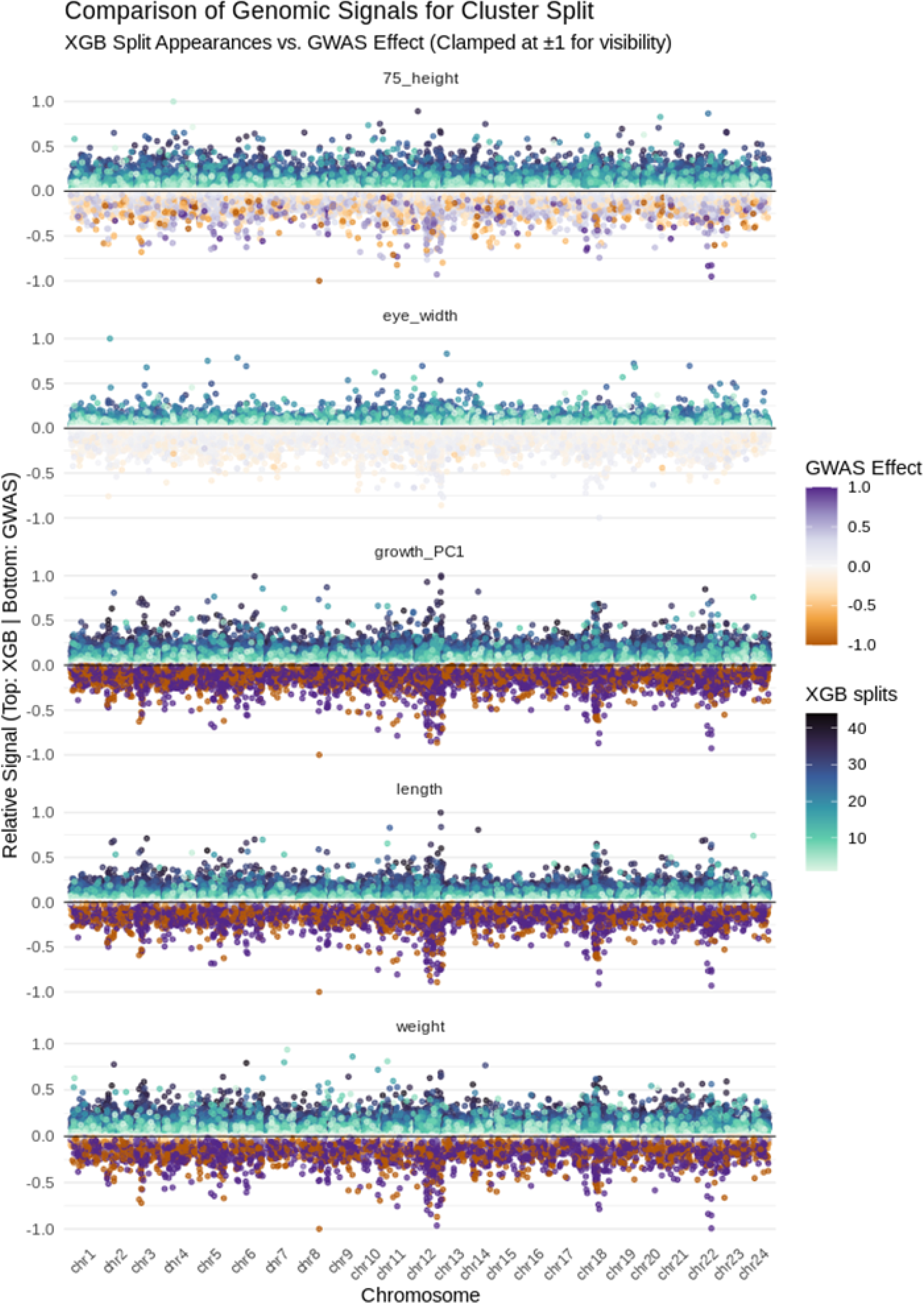
Miami plot contrasting XGBoost feature stability and GWAS effects across growth phenotypes under the genetic cluster validation split. Each horizontal track represents an individual morphometric trait. Positions above the zero line track XGBoost scaled feature importance (gain), with points color-coded by the number of cross-validation splits in which the SNP was retained. Positions mirrored below the zero line track GLM GWAS significance, color-coded by estimated marker effect size (purple-to-orange scale). Extreme GWAS effect values and XGBoost gain are squished at +/-1 to preserve background visual resolution and facilitate direct peak comparison between machine learning selection and traditional association mapping.

### Genomic prediction using GBLUP

GBLUP predictions were more accurate (higher R^2^) for the training set than testing set in all traits predicted. The mean training R^2^ across 50 CVs for the random individual split ranged from 0.385 (eye_width_mm) to 0.575 (weight), but for the testing set this dropped to a range of 0.085 (eye_width_mm) to 0.146 (weight). For the relatedness aware clustering, in the training set the R^2^ values ranged from 0.389 (eye_width_mm) to 0.581 (weight) and 0.055 (length) to 0.075 (weight) in the testing set (Figure 4A, Supplementary Table 4).

**Figure 4.**
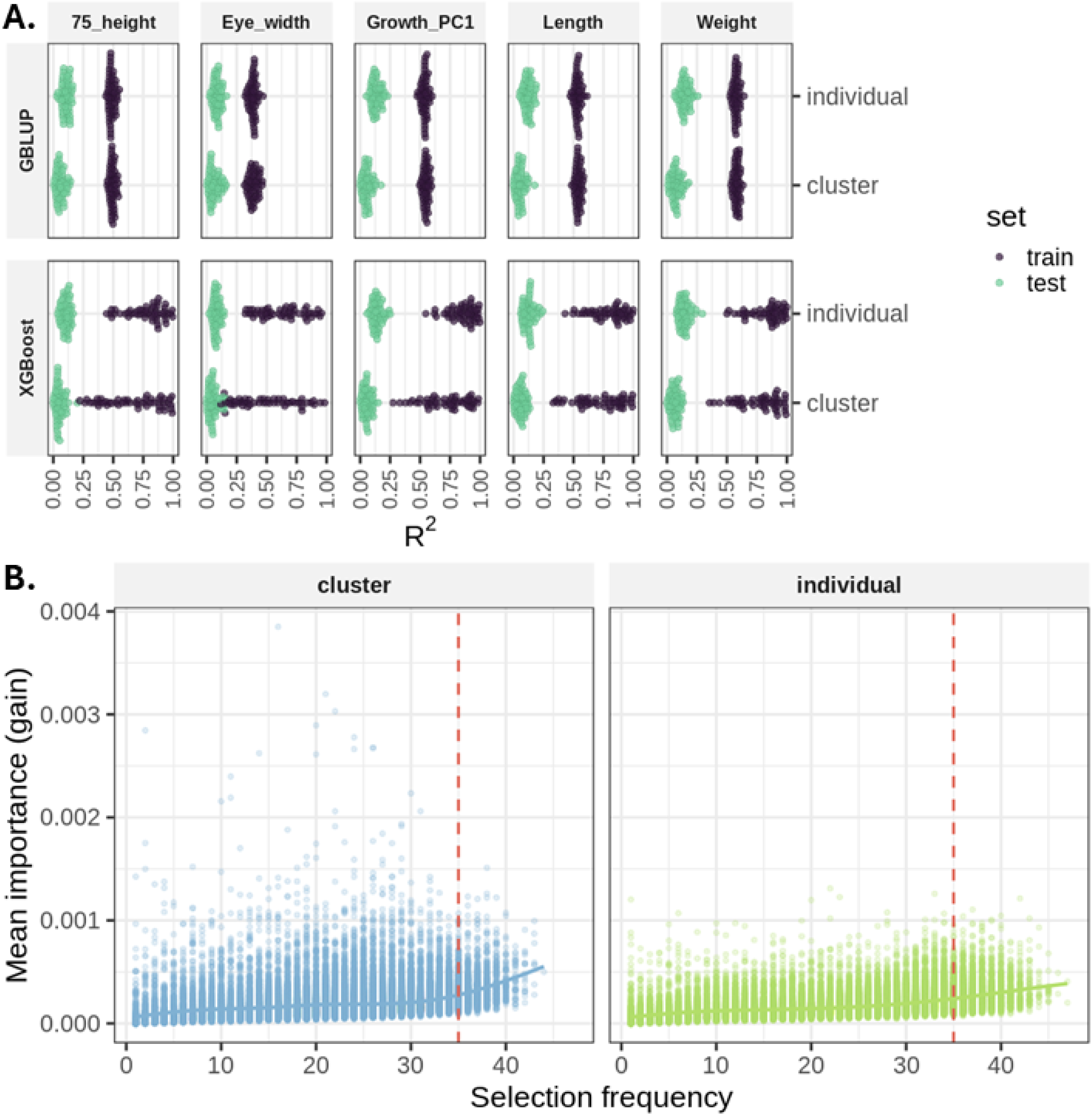
**A)** Beeswarm plots showing the R^2^ values of individual cross validations (CVs) for GBLUP (top) and XGBoost (bottom). Each plot shows the results for training (purple) and test (teal) data sets, as well as the data stratification approach (individual = random split, cluster = relatedness aware stratification). **B)** The mean importance (gain) of each SNP selected by XGBoost and the number of CVs each one appears in. The relatedness aware data stratification approach is on the left, and random split on the right. The dashed red line shows the 35 CV threshold for persistently selected SNPs.

### Genomic prediction and identifying growth-related SNPs using machine learning

There were two approaches taken for XGBoost, 1) a random train (80%) test (20%) split of individuals and 2) a random train (80%) test (20%) split of entire relatedness clusters. The mean R^2^ values for the training set across the 50 CVs in the first approach ranged from 0.631 (eye_width_mm) to 0.855 (PC1 of traits), while the testing set from the same approach had mean R^2^ values between 0.079 (eye_width_mm) and 0.145 (weight). The relatedness aware stratification approach resulted in mean training R^2^ ranging from 0.504 (eye_width_mm) to 0.767 (weight), while it ranged between 0.052 (height at 75% of the fish) and 0.066 (weight) in the testing set (Figure 4A, Supplementary Table 4). Across the 50 cross-fold validations, 10,840 SNPs appeared in at least one feature importance matrix for the individual-based split, and 10,816 SNPs for the relatedness aware split. The mean gain per SNP across the 50 splits corresponded with the GWAS p-value peaks (Figure 3 asnd Supplementary Figure 2). A subset of the ∼10,000 SNPs selected as important features in any given XGBoost CV (random split = 3,413; relatedness aware split = 1,786) were present in 35 or more splits for any given phenotype (Figure 4B, Supplementary Figures 3 and 4). Of these, 730 persistent SNPs in the relatedness aware data split approach were shared across more than one phenotypic measure, and 1450 persistent SNPs in the random split approach were shared across more than one phenotype (Supplementary Figure 4). In both test:train split approaches, the mean gain of persistent SNPs (≥35 CVs) was a significantly greater than that of background SNPs (< 35 CVs, Mann-Whitney *U* test, p < 0.001; Figure 4B, Supplementary Figure 3). Under the random test:train split, the mean gain of the persistent SNPs was 0.000264 (95% CI: [0.000261:0.000268]), nearly twice that of the background SNPs (mean gain: 0.000163, 95% CI:[0.000162:0.000163]). Similarly, the relatedness aware test:train split resulted in a mean gain of persistent SNPs of 0.000312 (95% CI: [0.000306:0.000318]) compared to 0.000180 for the background SNPs (95% CI [0.000179:0.000181]).

## Discussion

This study demonstrates how high-throughput phenotyping, genomic prediction, genome-wide association analyses, and machine-learning approaches can be integrated to support selective breeding programmes. Using snapper as a case study, an emerging species for aquaculture, we show how image-based phenotyping can generate scalable measurements of growth and morphology, which can then be linked with genomic information to estimate breeding values, identify trait-associated variants, and explore alternative prediction frameworks. One of our findings is that image-based phenotyping not only increased the number of measurements collected from individuals but also resulted in the discovery of more significant SNPs via GWAS. This was not only because more measurements and traits could be analysed via GWAS, but derivatives of those allometric and complex traits could also be more fully explored and subsequently included if needed. We were also able to predict weight using both GBLUP and XGBoost, capturing between 18% and 39% of total heritable variance (h^2^ = 0.38). While both models showed high training accuracy, the observed overfitting, especially in the relatedness aware based data stratification approach, likely reflects the reproductive skew observed in this species (Ashton et al. 2019), and the estimated low effective population size of this cohort. Despite these challenges within this dataset, XGBoost identified similar biological signals to traditional linear methods, highlighting shared genomic hotspots for growth.

Computer vision methods are becoming increasingly common in biology for tasks including species identification, population census counts, and phenotyping (Yang et al. 2021; Fu and Yuna 2022; Tuckey et al. 2022; Babu et al. 2023). Here, we were able to increase the number of measurements taken from each individual while decreasing the amount of labour involved and stress on the fish (due to a reduced time out of the water). Image-based, computer vision phenotyping could also be extended to underwater photographs or videos in the future, further minimising work and welfare impacts and increasing data volume and potentially enabling real time assessments (Fu and Yuna 2022). The increased data in this instance allowed for the identification of 23 additional growth-associated SNPs via GWAS compared to only analysing weight and length. A similar approach has already helped similar analyses aiming to uncover the genomic architecture of growth or other traits of interest in mungbean (Boddepalli et al. 2025). Growth is particularly suited for computer vision phenotyping since many growth-related traits can be collected from images. Additionally, the strong allometry of these measurements can enhance statistical power by better partitioning phenotypic variance or providing avenues to reduce dimensionality. Combined, this can help understand the genetic complexity of such a polygenic trait. This may be particularly helpful in species where resources don’t allow for large sample sizes and genetic panels such as in humans.

The efficacy of genomic prediction and trait discovery are not just limited by phenotypic data availability, but experimental design artefacts, population genetic factors and genetic relatedness can influence these analyses too. In our analyses, the resolution of the genetic signals was also limited by the reduction of estimated genomic *N_e_* in our breeding programme cohort, from ∼117-176 in the founding population in Tasman Bay (Hauser et al. 2002) to 18.2-57.8 in the cohort included here. While using the interchromosomal LD interactions mitigates the confounding effects of physical linkage, N_e_ estimates calculated in this way can still be driven down where populations do not meet Wright-Fisher assumptions (Waples 2025). Within the context of this cohort of snapper, there has been a history of strong recent selection for growth traits, a high bias in parental contributions where individual males contribute more offspring than individual females, and a recent reintroduction of wildtype fish to mitigate inbreeding (Ashton, Ritchie, et al. 2019; Ashton, Jaksons, et al. 2019; Samuels et al. 2024). These factors together likely explain the very few strong interchromosomal LD interactions through epistatic interactions for selected traits and incomplete dissipation of LD from the recent introgression of wild broodstock. Similar strong interchromosomal LD patterns have been attributed to selection and epitasis in other species (e.g (Tessele et al. 2025)). Typically, a smaller *N_e_* would result in more accurate predictions (Goddard 2009). However, the factors outlined above that may explain the depressed N_e_ estimates in this cohort would also likely increase overfitting in genomic prediction.

Such a relatedness structure in the F_4_ cohort of these snapper would no doubt lead to higher data leakage and causes overfitting to be amplified. Data leakage was indeed more apparent in the random split approach to data stratification because more of the siblings and half-siblings of the training set fish were in the testing set. This allowed the model to be informed by related individuals rather than broad biological signals. The disruption of this leakage in the relatedness aware data stratification did lead to a reduction in over-fitting, but also in the prediction accuracy. Nonetheless, some (∼18%) of the genetic heritability for the traits was still able to be captured by the model when controlling for relatedness. This is lower than that captured in another study comparing genomic prediction in other aquaculture species, but our results from the random split are more similar, and data leakage control due to relatedness does not seem to have been considered (Kriaridou et al. 2020). Our second approach highlights the challenges of capturing generalisable genomic signal in a genetically narrow cohort of a breeding programme, especially for a complex polygenic trait, regardless of whether linear or non-linear frameworks were used. This limitation of genomic prediction, where narrow genetic bases reduce the resolution of causal variant detection and trait prediction, is widely documented across breeding programs and evolutionary studies (Wellenreuther and Hansson 2016; Song et al. 2023; Yáñez et al. 2023). Consequently, both traditional linear models and more complex non-linear machine learning frameworks approach the same performance ceilings when population diversity is constrained (Chafai et al. 2023). Despite this, genomic prediction can still be meaningful in aquaculture programs through reduction in breeding interval, balancing strict selection with genetic diversity maintenance, and when applied to phenotypes that are difficult to assess (Chafai et al. 2023; Yáñez et al. 2023).

Beyond the limitations of population structure in our dataset, the performance of different predictive frameworks suggests that the genetic architecture of growth in this cohort of snapper seems to be primarily driven by additive effects. Both GBLUP (linear) and XGBoost (non-linear decision tree) arrived at similar R^2^ values in the test sets (∼0.145 in randomly split data and ∼0.075 for relatedness-based data stratification), and XGBoost identified similar genetic signals to GWAS (also linear). While the GLM’s overinflated p-values captured a mix of true biological associations and confounding population stratification, it provided a valuable continuous baseline of regional genomic signal. By plotting XGBoost features gain against the GLM (Figure 3), we visually evaluated how effectively our downstream cross-validation frameworks purge this background inflation. SNPs that track with inflated GLM peaks but were discarded by the relatedness aware stratification confirm the model’s ability to identify and strip away population-specific confounding noise. Conversely, regions where robust XGBoost features persisted across population clusters (i.e. coinciding with prominent GLM peaks but missed by the conservative FarmCPU multi-locus penalty) highlight candidate loci where non-linear architectures or complex local LD blocks may have been rescued by machine learning. Polygenic traits such as weight often have a non-additive, epistatic underlying genetic architecture that is not captured by linear model analyses (Wellenreuther and Hansson 2016; Ruigrok et al. 2022; Yengo et al. 2022; Križanac et al. 2025). Non-linear models such as decision trees can capture these kinds of genetic interactions. If such architecture was present in a dataset, the training and testing set performance using a tool such as XGBoost should be higher than that of GBLUP or GWAS. However, our results only showed a higher accuracy for XGBoost in the training set, but not testing set, suggesting either that there is no non-additive signal in the genetics of growth of this cohort of fish, or our sample size and data structure prevented its detection. Simulations and empirical analyses have shown that additive linear models often absorb substantial non-additive effects (Morgante et al. 2018). However, as the number of QTLs associated with a trait increases, non-linear methods tend to lose their predictive advantage over linear frameworks in the context of epistasis (Daetwyler et al. 2010). Application of these methods in larger datasets without the genetic structure in ours or to less complex traits could better test the ability of XGBoost to identify these non-additive effects. The concordance of the linear and non-linear methods, especially the identification of overlapping genomic hotspots for growth related variants, does confirm the robustness of the application of decision trees to this genomic prediction problem.

## Conclusions

This study has demonstrated how high-throughput, image-based phenotyping can be paired with genomic prediction methods to understand genomic architecture of complex traits and guide breeding decisions in an emerging aquaculture species, the Australasian snapper. The implementation of computer vision phenotyping expanded the data volume and increased the number of SNPs discovered by GWAS, but genomic prediction was hampered by data leakage and population structure stemming from reproductive bias and the design of the breeding program. While the non-linear prediction method (XGBoost) did not outperform the linear approach of GBLUP in this instance, XGBoost did identify genomic signals overlapping to those identified using the linear GWAS approach, and resulted in a larger list of potentially important variants. This type of finding should be validated in species with broader functional genomic resources. Ultimately, our findings highlight the utility of machine learning in phenotyping and genomic prediction, even if prediction accuracy faces the same hurdles as those faced by GBLUP.

## Declarations

### Ethics approval and consent to participate

All work was done under the conditions of animal ethics application AEC-2021-PFR-05, approved by the Animal Ethics Committee at the Nelson Marlborough Institute of Technology (Te Pūkenga), along with fish-farm licence FW208.03.

### Consent for publication

Not applicable

### Availability of data and materials

The genotyping data used in this work are already published (Montanari et al. 2023) as is the genome assembly (Blommaert et al. 2024). Both these datasets are available with permission from representatives of Māori iwi (tribes). Guardianship of these datasets are managed by the Aotearoa Genomic Data Repository (Te Aika et al. 2023).

## Authors’ contributions

JB analysed and interpreted the data and was a major contributor to writing the manuscript. PEB analysed and interpreted the data and contributed to writing the manuscript. DA and GS provided the pipeline for analysing the phenotypic data. LJ contributed to data analysis, writing and editing. MW contributed to conceptualisation, funding acquisition, data interpretation, writing, editing, and overall manuscript development. All authors read and approved the final manuscript.

## Acknowledgements

We thank all the staff of The New Zealand Institute for Plant and Food Research Limited who were involved in breeding and rearing the snapper populations that formed part of this breeding programme.

---

**Supplementary Figure 1.**
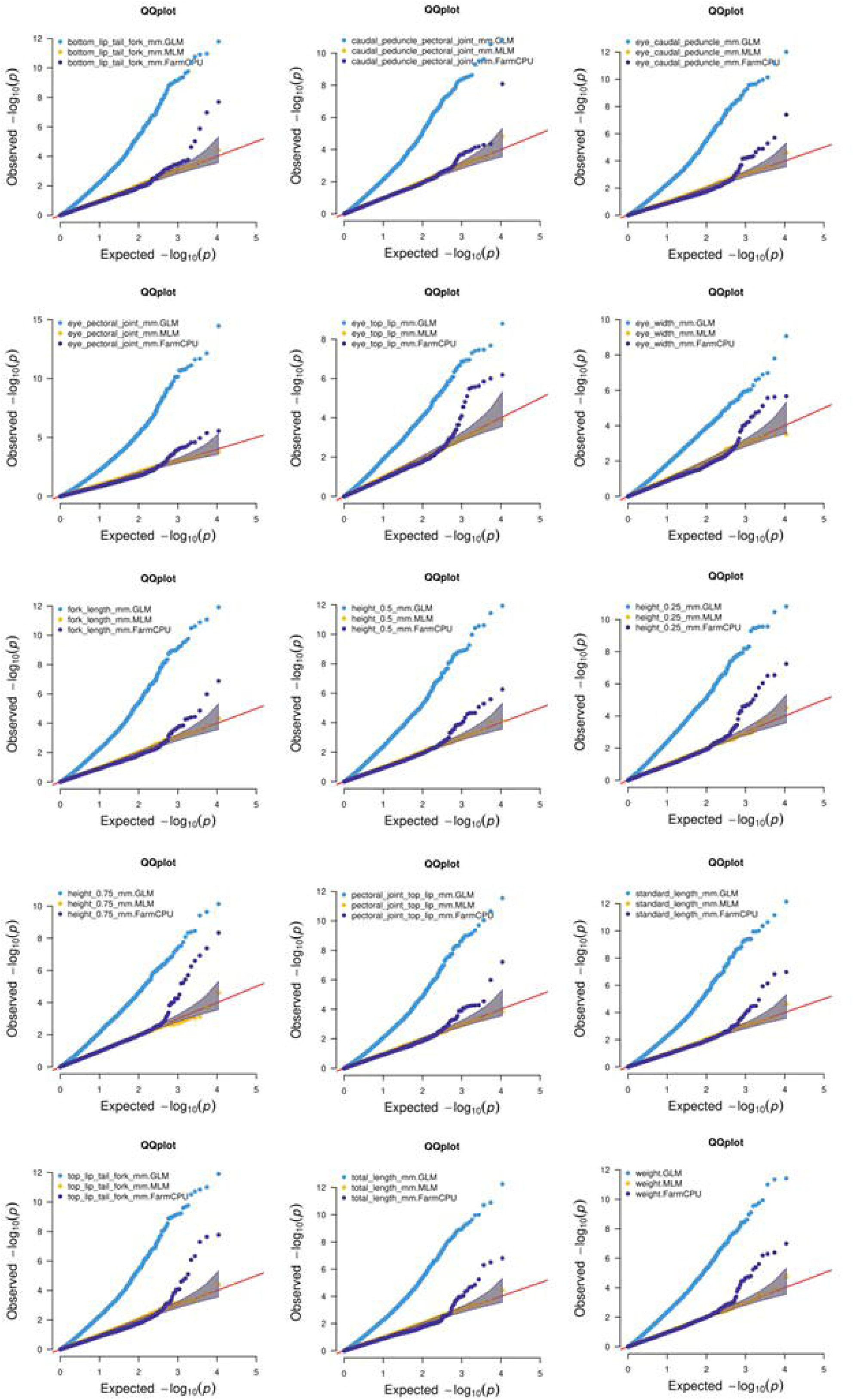
Quantile-quantile plots showing the observed versus expected – log_10_(p) values for the three methods (GLM, MLM, FarmCPU) for all measurements.

**Supplementary Figure 2.**
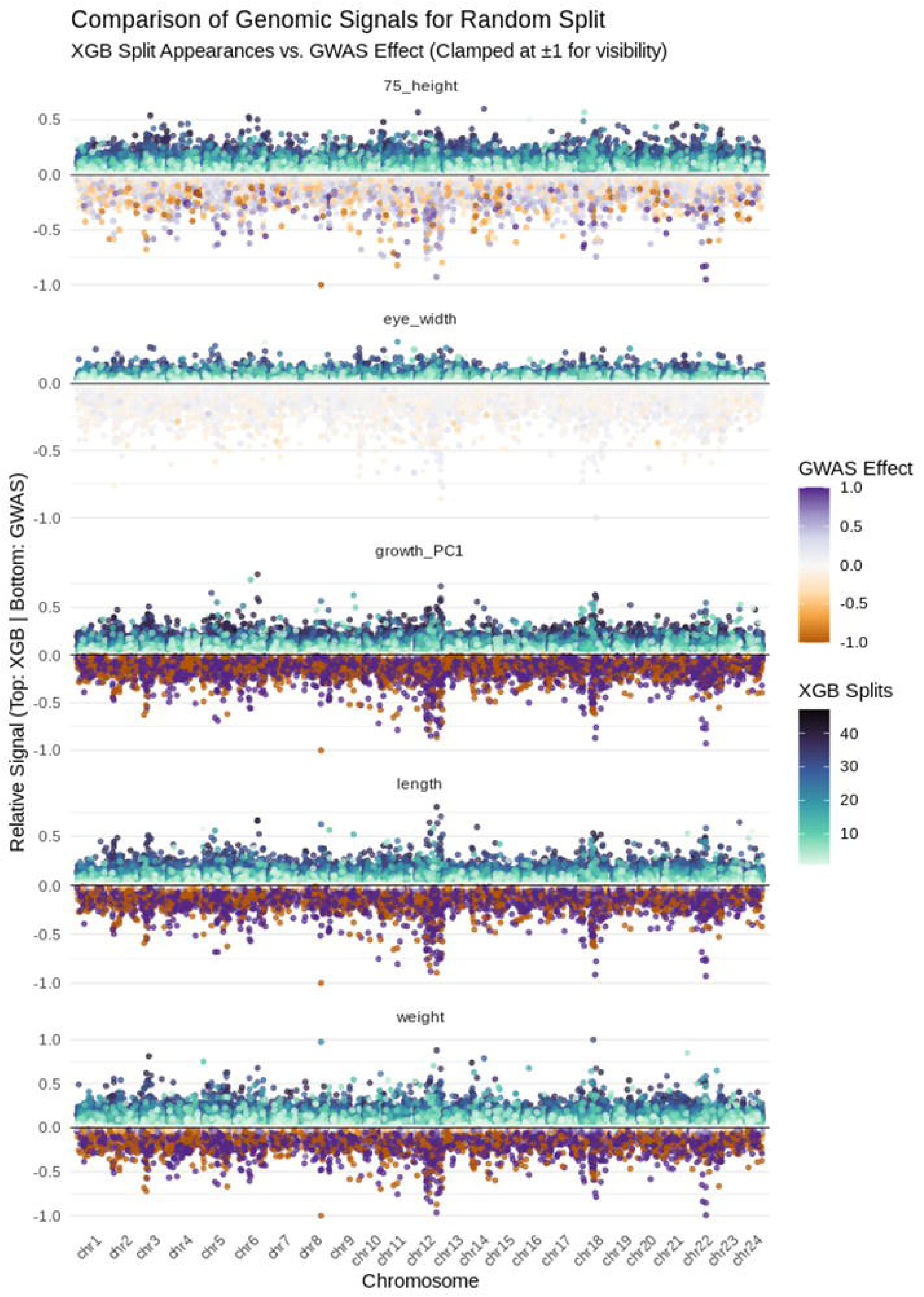
Miami plot contrasting XGBoost feature stability and GWAS effects across growth phenotypes under the relatedness cluster validation splits. Each horizontal track represents an individual morphometric trait. Positions above the zero line track mean XGBoost scaled feature importance (gain), with points color-coded by the number of cross-validation splits in which the SNP was retained. Positions mirrored below the zero line track GLM GWAS significance, color-coded by estimated marker effect size (purple-to-orange scale). Extreme GWAS effect values and XGBoost gain are squished at +/-1 to preserve background visual resolution and facilitate direct peak comparison between machine learning selection and traditional association mapping.

**Supplementary Figure 3.**
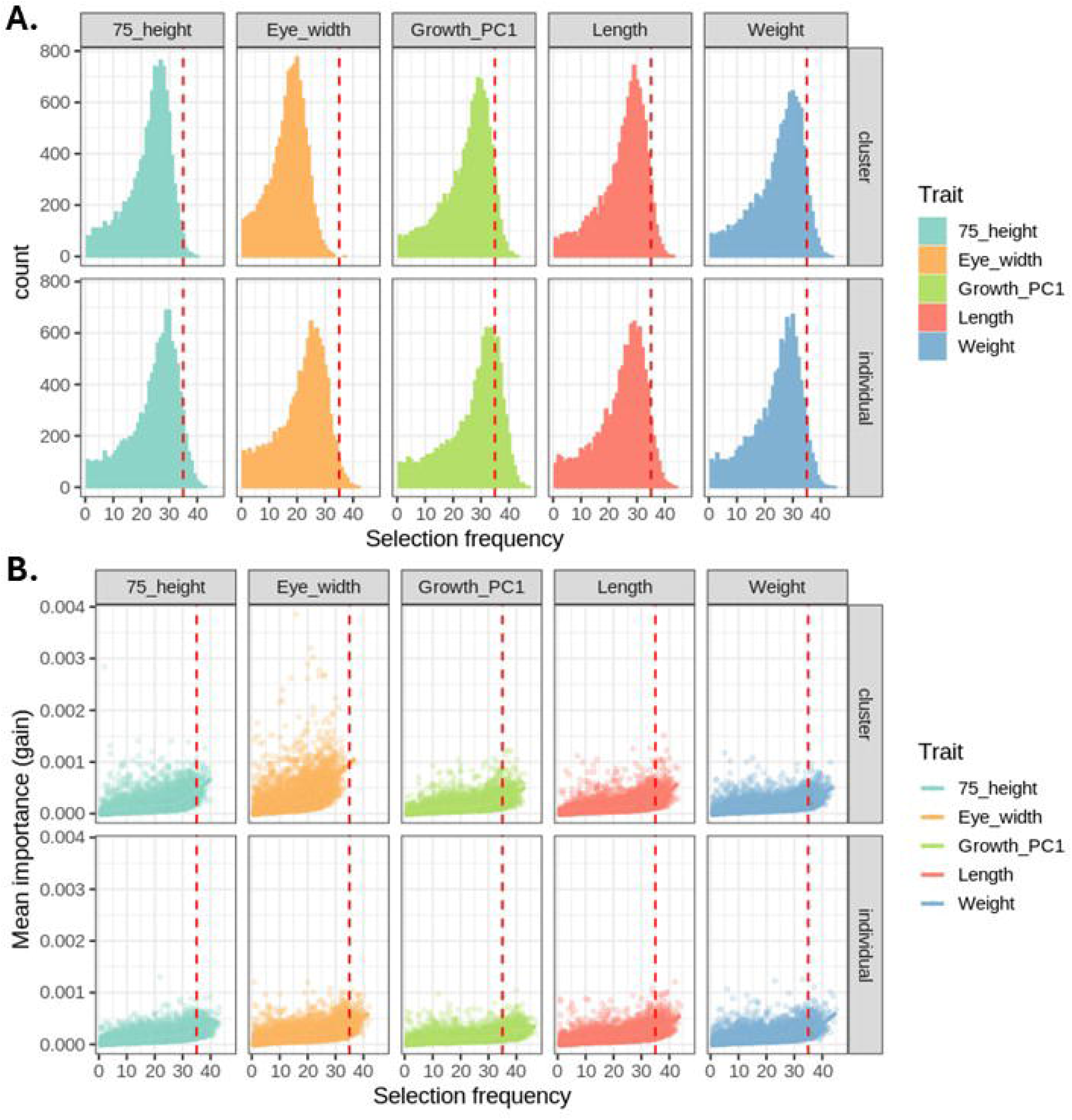
**A)** Histogram of the number of CVs (Selection frequency) a SNP appears in an XGBoost feature importance list for each data stratification type and phenotype. **B)** The mean gain for each SNP and number of CVs (selection frequency) a SNP appears in a feature importance list in XGBoost for each data stratification type and phenotype. The red vertical line in both plots indicates a threshold of 35 CVs to define persistent SNPs.

**Supplementary Figure 4.**
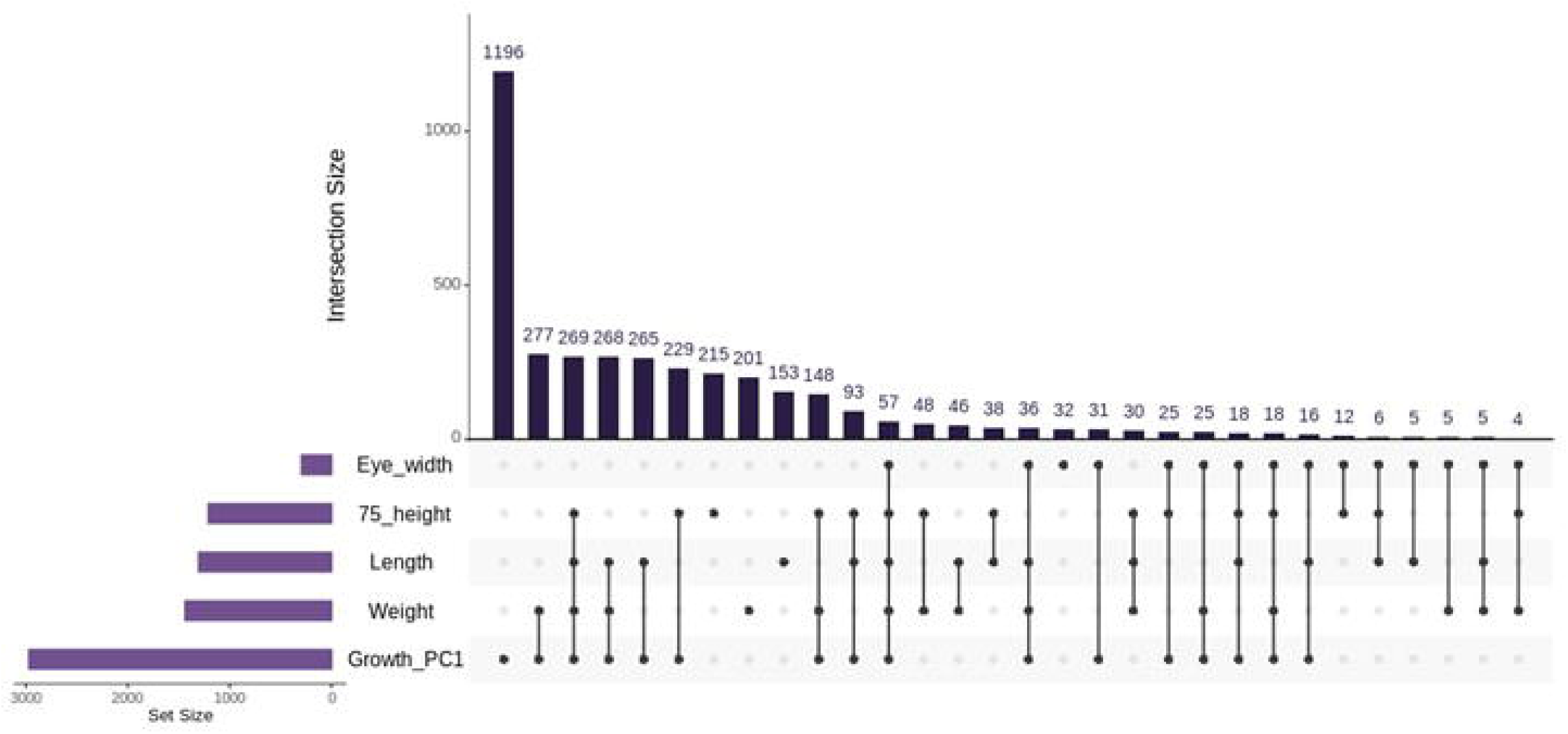
An upset plot showing the overlap of persistent SNPs across the 5 phenotypes analysed. Horizontal bars (left) indicate the number of persistent SNPs per phenotype (set size), filled dots connected by lines indicate which overlap of SNPs is being shown by the black bar which indicates the intersection size in the main plot.

